# Growth, survival, and fitness in the first year of life for *Pycnopodia helianthoides* under different larval densities

**DOI:** 10.64898/2026.04.02.716152

**Authors:** Elora H. López-Nandam, Logan Tallulah Story, Mariah Evin, Jessica Witherly, Margarita Upton, Lana Krol, Freeland Dunker, Kylie Lev

## Abstract

Sea star wasting disease has caused widespread mortality in the kelp forest predator, the sunflower sea star (*Pycnopodia helianthoides*). Wild populations have declined by up to 99% in parts of their native range along the western North American coast. In response, a multi-institutional conservation breeding and rearing program has been initiated to support future reintroduction efforts for the species. We split a full-sibling cohort across four larval density treatments (1 larva/ml, 2 larvae/ml, 5 larvae/ml, and 15-20 larvae/ml) to assess the effects on larval settlement, juvenile survival, and juvenile fitness at 12 months old. Stars raised in the highest density treatment displayed a lower settlement rate and were significantly smaller than the other density groups at 12 months old, but showed no significant difference in flip time, a measure of fitness. Additionally, measurements of diameter, weight, and arm count across modern and historical juvenile and adult stars indicate that *P. helianthoides* experience exponential weight gain as they grow in length, with corresponding asymptotic growth in arm count. These findings will inform best practices for the aquarium propagation of *P. helianthoides* and will contribute to broader efforts aimed at reestablishing populations in the wild.

## Introduction

In 2013, a mass mortality event caused by sea star wasting disease (SSWD) affected over 20 species of sea star along the western North American coast, from Baja California through Alaska, resulting in the mortality of billions of sea stars [1–3]. The largest-bodied of these species, the sunflower sea star or *Pycnopodia helianthoides*, (Brandt 1835) was the species with the highest mortality rate. The southern end of the range (Baja California and California) experienced 99% or higher mortality [4]. Sunflower sea stars are keystone generalist predators on the kelp forest floor, whose diet includes a variety of gastropods, bivalves, crustaceans, and other echinoderms, including sea urchins and sea stars [5,6]. In the years following the SSWD outbreak, kelp forest cover has dramatically declined in large part due to overproliferation of purple sea urchins. Some have hypothesized that the increase in purple sea urchin population sizes and consequent overeating of kelp are related to the near-disappearance of their once-predators, the sunflower sea stars [7]. There have, therefore, been increasing calls for sunflower sea star breeding and reintroduction as one possible approach to regenerating balance in kelp forest ecosystems [8]

Echinoderm spawning and larval culturing have been studied since 1847, historically as model systems for embryological studies [9]. Raising these animals out through settlement and then the full life cycle, however, is more recent. For some marine invertebrates, population density has declined to the point where natural recovery is unlikely due to Allee effects [10]. The aim of burgeoning conservation breeding efforts is to raise healthy, competent animals that can be released in the ocean, thereby increasing population density and genetic diversity, so that populations can rebound [11]. Recognition of the ecological importance of sunflower sea stars, and recent advances in spawning and rearing these animals in human care [12,13], have led to great interest in raising these animals at scale.

When considering scaling a marine invertebrate breeding program, larval density is among the first aspects to be considered. Increasing the density of larvae in a container of seawater reduces the amount of space needed to keep a large number of larvae, but it may change the water quality (oxygen, pH, ammonia, organic matter) of the system. Increasing larval density may also result in increased stress or competition for space or food, which in turn may lead to tradeoffs in mortality, size, or fitness of the larvae, or later, of the juveniles [14]. While some studies have examined the effects of larval density on larval mortality rate and growth in echinoderms [14–17], these studies all ended prior to or just after metamorphosis, rather than follow the effects on the resulting juveniles well after settlement. The effects of larval density on larval size, growth, and survival are not consistent across these studies, although that may be confounded by the fact that these studies all looked at different species. Considering that the environmental conditions experienced by larvae can influence their performance in later life stages [18–20], more work following sea stars raised at different densities to later in life is needed in order to inform decisions made by conservation breeding programs.

Here, we present the results of spawn induction via 1-methyladenine (1-MeAde) of one male and one female sunflower sea star, and the subsequent raising of the cohort of full siblings produced by this mating. We followed the time to and rates of settlement, growth, survival, and fitness of the cohort raised at four different larval densities (1 larva/ml, 2 larvae/ml, 5 larvae/ml and 15-20 larvae/ml) for one year post-fertilization. We also combined our one year-old juvenile size measurements with datasets of live and preserved adult sunflower sea stars, to model the relationships between number of arms, length, and weight as sunflower sea stars grow.

## Materials and methods

### Spawn induction and fertilization

Spawning was induced in two adult sunflower sea stars held at Birch Aquarium in San Diego, CA, on February 14, 2024. The resulting spawn, larvae, and juveniles are referred to as the Cupid Cohort. Prior to spawn induction, one male and one female sunflower sea star were pulled from their exhibit to be weighed and placed into separate flow-through strainers in a water bath. Each adult was injected with 1-MeAde, an aminopurine compound produced by the follicle cells of the sea star that forms a component of the spawning response [21,22]. When exogenous 1-MeAde is administered to a sea star, the follicle cells separate from the oocyte surface, ultimately resulting in spawning. For full spawn induction protocol, see Supplementary Methods.

Once spawning commenced, 5-gallon buckets with monolayers of eggs were set up in the water bath and diluted sperm was added. To dilute the sperm, we used one drop of concentrated sperm per 15 mL of filtered sea water and mixed gently until the sample looked milky and completely combined. 15 mL of the sperm dilution was added to each 5-gallon bucket of eggs. We fertilized 7 buckets with fresh sperm. We waited 10 minutes and then checked for fertilization via microscope. Once fertilization was confirmed, the fertilized eggs were rinsed of excess sperm and topped off with filtered seawater for an overnight water bath.

The 5-gallon buckets with banjo filters were left in the water bath overnight. The next morning the larvae were transferred from the plastic 5-gallon buckets into 1 L glass transport jars, which were placed in a cooler and secured with styrofoam. The temperature was controlled with ice packs. Other embryos were transferred to other institutions for rear out.

### Culture set up

Sunflower sea stars and echinoderm food cultures were reared at the Steinhart Aquarium utilizing two 3’x4’ recirculating water tables. Each system has a total running volume of approximately 225 gallons utilizing artificial seawater made in-house. To maintain a temperature of 12.8° C, all cultures were partially submerged on the table from 2-6” at all times. Water changes in standing-water cultures (jars) utilized water directly from the tables for water changes, otherwise cultures were recirculating. Water filtration consists of an MRC Orca Pro II fractionator, Aqualogic heat exchanger, UV sterilizer, bio-tower, and a 1µm filter. Lighting consisted of (4) Kessil A500X set at approximately 75% intensity with a photoperiod of 10 hours light/14 hours dark, positioned 36” above the cultures to maintain an even light distribution.

Water quality parameters were maintained with a pH 8.1-8.3, salinity 33-35 ppt, temperature 12.2-13.3 °C, Ammonia 0 mg/L, Nitrite <0.2 mg/L, Nitrate <10 mg/L, Alkalinity range of 3-3.5 mEq/L, and calcium range of 400-430 mg/L.

### Larval rearing

Larvae were reared in 8 L glass jars or 15 L plastic cambro buckets. Larvae were diluted to 1 larva/mL (n = 8), 2 larva/mL (n = 4), and 5 larvae/mL (n = 4) in the glass jars. All remaining larvae were evenly distributed among the plastic buckets at 15-20 larvae/mL (n = 4).

StirMATE Variable Speed Smart Pot Stirrers (StirMATE, Orinda, CA, USA) were used to keep larvae suspended in the water column without damaging the larvae, as described in [23] (Supplementary Figure 1). We replaced the built-in stirring attachment with a silicone spatula. All silicone spatulas were angled to have the convex side pushing outwards and forward in the direction of the water flow. This helped to prevent damaging larvae. We extended the StirMATE arm so that it went directly over the middle of the container. The stirMates were set to move clockwise, with the spatula extending from the middle of the culture to the wall of the jar or bucket, without touching the wall. Flow of the StirMATE was adjusted to the nine o’clock position with the on dial. StirMATEs were always plugged in, rather than relying on battery power. They were checked twice daily to ensure that they were all operating properly.

Larval jars and buckets received water changes three times per week: two 50% water changes and one 100% water change with a jar change at this time. Jars were acid washed after the 100% water change. Water changes were done through a 120µm screen initially, and then through a 250µm screen once the majority of the larvae in the culture reached the brachiolaria stage.

### Larval measurements

Larvae were measured and photographed using an Olympus SZX10 microscope (Olympus Corporation, Tokyo, Japan) and the Infinity Analyze version 7 attachment. We measured larvae on a 1 mL sedgwick slide, which has a grid for better referencing for photos and measurements. Measurements were taken from the anterior to the posterior of the larva. Taking measurements from the median dorsal lobe to the rudiment for all larvae created the most uniform measurements. We measured 2-12 larvae per density treatment per timepoint to calculate average sizes (Supplementary Table 1).

### Feeding

Sunflower sea star larvae were supplied a diet of the phytoplankton *Rhodomonas lens* twice daily at approximately 9 am and 4 pm. For details on phytoplankton culturing, see Supplementary Methods. Ten individuals from each jar were regularly subsampled and their gut tracts were examined visually under a microscope to determine gut content and potential satiety. If larvae sub-sampled in a culture were found to have empty digestive systems before their 9 am feed, then food was increased until sub-sampled larvae retained food throughout the night. This provided the larvae with adequate nutrition while maintaining the cleanliness of the cultures by avoiding potential overfeeding.

Juvenile (post-settlement) stars were fed either live or frozen food. Of the live foods, purple urchins (*Strongylocentrotus purpuratus*), bat stars (*Patiria miniata*), and ochre stars (*Pisaster ochraceus*) were reared in-house. Bat and ochre star spawn induction followed the same protocol as the sunflower sea star 1-MeAde induction described above, with the exception that sperm was slowly added to eggs, eggs were gently agitated with a glass rod, and this was repeated until fertilization rate reached 90%. Larval bat and ochre stars were kept in the same glass jars,rigged with StirMATEs, and also received a diet of *R. lens*.

Purple sea urchins weighing 150-300 g were injected with 0.75mL of 1M KCl. Once sperm and eggs were produced by the urchins, the fertilization and rearing process was identical to that described above for the bat and ochre stars. Cultures that were kept without direct light did not develop correctly, and larvae were found dead on the bottom of their culture vessels. The lighting described above under *Culture set up* allowed for successful larval rearing.

Other live foods offered to settled stars included tropical tuxedo urchins, asterina sea stars, spirobis worms, softshell clams, and oysters. They were also offered 150-250 µm otohime, frozen cyclops, calanus, and finely grated LRS Reef Frenzy (LRS Foods, North Carolina, USA). As juveniles grew, the following food items were added and cut to appropriate size (less than ½ of the star’s body length) for the animals: pacifica and superba krill, hikari mysis, prawn, squid, clam tongue, mazuri carnivore and low fat gel diets. Capelin and mackerel were also offered, but routinely refused.

### Juvenile rearing

Metamorphosis and settlement occurred in larval vessels without addition of settlement cues. Settled stars were counted (Supplementary Table 2) and then a soft bristled artist’s paintbrush was used to gently move the newly settled stars into settlement bins. Settlement bins were originally white plastic bins measuring 11” x 14” x 5”. All four upright walls had windows with 100 micron screens, to allow water flow. White bins were replaced with black bins two months post settlement, as the settled stars were easier to see and track against the darker background.

Until the sunflower sea stars were strong enough to cling tightly to the settlement bins, we cleaned settlement bins by rinsing bin inhabitants into a sieve, and attempted to flush the unwanted material out of the sieve without damaging the stars. Once the stars were large enough to cling to the bins, we could begin siphoning unwanted material from the bins and fewer outbreaks of copepods and “pink slime”—a pinkish red biofilm that occasionally formed in the bins and caused mortality in any stars that touched it. The slime biofilm was treated by carefully moving all remaining stars with a paintbrush into a clean bin and then sterilizing the infected bin by soaking it in bleach (10 mL of 8% bleach per 1 gallon of water) and then neutralizing with sodium thiosulfate.

### “Monster” star separation

In all settlement bins, one or several stars increased in size more rapidly than their siblings. These larger stars were frequently observed feeding on their siblings, hence dubbed “monster stars”. To reduce cannibalism and level the competition’s field, these large stars were removed from the communal settlement bin and placed into solitary bins measuring 3.5” x 2.5” x 2.5”, referred to as “condos.” They were then moved into larger bins made out of modified 1 L plastic bottles to allow space for growth seven months post settlement. It was noted that animals kept in large containers grew to large sizes whereas animals remaining in the smaller condos remained small.

All larval containers were given a name and tracked through time in settlement bins and condos so information on initial larval densities, larval vessel types, and results from the fitness tests could be tracked.

### Fitness tests

At one-year post-fertilization, juvenile stars went through a series of “fitness tests.” Data recorded during the fitness test included a righting (or flip) trial, and recording the weight, number of arms, and total length of the animal (Supplementary Table 3). A secondary test was conducted to see the stars’ response to food as well.

The righting trial (referred to as the flip test) was performed by placing the animal onto its dorsal surface and timing how long each animal took to right itself to its original position—ventral side attached to substrate. Times were recorded to the approximate second when a full flip was achieved by the individual. The timer was stopped and the test ended if the star could not right itself after 30 minutes had elapsed.

To capture juvenile star weight, individuals were placed in a small bin filled with artificial seawater. Prior to weighing, the water and the bin were zeroed for accuracy. Weight was recorded in grams, and the same scale and bin were used for all individuals. The arm count was recorded at this time, counted as the number of limbs on each individual from the dorsal side.

Length was captured as the longest armtip to armtip measurement (Supplementary Table 4). It is important to note that measurements of sunflower stars can be highly variable based on positioning of the animal when measuring.

### Alaska adult star measurements

During a separate spawn event in 2025, the weight, length, and number of arms was measured for adult sunflower sea stars collected near Seward, Alaska, USA. When animals were removed from a habitat to be weighed, they were removed from the water and set upside down in a tray on a scale. While out of the water and in this position, the total number of arms were counted and the length of the stars was also measured when the stars were upside down using a cloth sewing measuring tape. Animals were also measured across the longest armtip to armtip diameter (Supplementary Table 4).

### Historical star measurements

Dry (dehydrated) and wet (preserved in 70% ethanol) sunflower sea star specimens held in the California Academy of Sciences (CAS) Invertebrate Zoology collection were measured for total length using the same methodology as both live specimens from Seward and the juveniles in the Cupid Cohort. Specimens also had their arms counted. The specimens measured were collected from across the Western Pacific (Gulf of Alaska, AK to Monterey Bay, CA) ranging from the years 1889 to 2023. To account for specimens that were stiff, inflexible, or fragile as a result of age or preservation, a string was marked from point-to-point of the arms then measured on a flat surface. Arm counts were determined by the number of clear, undamaged limbs or undamaged, newly developing limbs (≥ 2cm). Specimen measurements are enumerated in Supplementary Table 4, and are available online with photos (https://specify-portal.calacademy.org/iz/).

### Statistical analyses

Observations, measurements, and data throughout the larval and juvenile rearing stages were initially recorded in Google Sheets and then later exported to Microsoft Excel (Version 16.103.1). Individual data sheets were then saved as .csv files and imported into the R and RStudio (ver 4.5.2) statistical computing environment. To account for missing observations and random characters, data cleaning and filtering was performed using dyplr and tidyr packages [24,25]. Prior to statistical testing, assumptions of normality and homogeneity of variance were evaluated using a Shapiro-Wilk test and a Levene’s Test for Equality of Variances using functions from the car package [26]. Normality was assessed for each response variable using a significance threshold of α = 0.05. Data that did not significantly deviate from normality or homogeneity (p ≥ 0.05) were analyzed using parametric methods, whereas data exhibiting significant departures from normality and homogeneity (p < 0.05) were analyzed using non-parametric approaches. All statistical analyses and post-hoc testing were performed using pipe-friendly functions from the rstatix package [27].

To evaluate differential cumulative larval settlement between larval densities over time, a General Additive Model (GAM) was employed using the mgcv package [28]. A Negative-Bionomial family accounted for overdispersion in the normalized settlement data, smoothed over days post fertilization (DPF) by density treatment. Model outputs were transformed using the marginaleffects package [29]. Given the differential number of replicates between larval density groups, cumulative settlement data was normalized by dividing the number of settlers in each bucket with the number of replicates in each density treatment.

To examine the relationship between individual size, larval density, and flip time, flip time (seconds) was analyzed as a continuous response variable using linear modelling approaches. Initially, the relationship between flip time and juvenile size was evaluated using a simple linear regression with flip time (seconds) as the response variable and individual length (mm) as the predictor. Then, testing whether the effect of body size on flip time differed among larval density treatments, a second model was fit including density as a potential interacting predictor with length to evaluate the differences in flip time. Models were compared between one another and to a null model (response ∼ 1) using an analysis of variance (ANOVA) to determine whether inclusion of the interaction term significantly improved the fit. Model performance was assessed using adjusted R^2^ values and residual standard error. Significance of predictors was evaluated using two-tailed t-tests and overall model fit using an Overall F-Test.

### Non-linear Allometric and Asymptotic Growth Models

The relationship between body length and arm number was modeled using a nonlinear least squares regression model. A self-starting asymptotic growth function (SSasymp) was fitted using the nls() function from the stats package (R Core Team). The model is equated as:

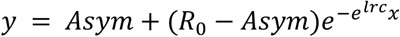

where *Asym* represents the horizontal asymptote of arm count, *R_0_* is the response at length zero, and *lrc* is the natural log of the rate constant. Model reliability was evaluated based on the number of iterations before convergence to the arm count asymptote.

To account for model prediction uncertainty, a nonparametric bootstrap resampling was performed. A total of 1,000 bootstrap replicates were generated by random sampling observations with replacement from the original dataset. The asymptotic model was refitted to each bootstrap using nls() and predictions were generated across 100 evenly spaced body length values spanning the observed size range. Model fitting errors during bootstrap iterations were handled using tryCatch(), and failed fits were excluded from subsequent analysis. For each length value, the mean predicted arm count and 95% confidence intervals were calculated from the bootstrap prediction distribution.

The model assumes that arm count initially increases exponentially with body length during early development, but eventually reaches an asymptote. Then a non-linear allometric growth model was employed, reflected as:

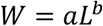

where *W* is body mass, *L* is body length, *a* is the normalized constant, and *b* is the allometric scaling component.

To linearize the model, both variables were log transformed, and a linear squares regression model was fit using the lm() function from the stats package (R Core Team). Parameter estimates were extracted from the fitted model and the intercept was then back transformed to obtain the models sampling coefficient *a*. Here, the slope of the linear regression is interpreted as corresponding to the allometric exponent *b*, with any deviation from *b* = 3 interpreted as evidence of non-isometric growth. Deviation from the allometric exponent would reflect changes in metabolic resource allocation as arm count asymptotes and, subsequently, weight increases. This model, similar to the non-linear asymptotic growth model, was run through a bootstrap sampling with the same technique in order to quantify uncertainty and perform robust checks on the model goodness-of-fit.

The nls() function and the non-linear allometric growth models did not natively compute a valid *R^2^*goodness-of-fit coefficient, so a *Psuedo-R^2^* statistic was calculated as:

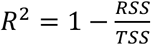

where RSS is the residual sum of squares and TSS is the total sum of squares relative to the mean of observed arm count, as normally calculated during regression analysis.

## Results

### Adult sunflower sea star spawning

The female sunflower sea star weighed 10.2058 kg and was given a 68.04 mL injection of 75 ppm 1-MeAde. The male weighed 11.3398 kg and received a 75.60 mL injection of 75 ppm 1-MeAde. The female injection started at 8:26 am and ended at 8:36 am. The male injection started at 9:00 am and ended at 9:09 am. We used two sticks per animal due to the limits of the syringe size and air getting into the IV extension when changing syringes. The female started spawning at 12:54 pm, 258 minutes after the last injection concluded. The male started spawning at 11:50 am, 161 minutes after the last injection concluded. Animals were monitored post spawn, and were confirmed healthy and resumed normal behavior and eating patterns within a day post spawn.

### Larval size over time and larval settlement by density treatment

Larval size increased over time across all density treatments from 4 to 47 days post fertilization (DPF) (Figure 2). Early in development, mean larval size when measured was similar across density treatments. However, divergence in growth trajectories became apparent at approximately 30 DPF, after which density-dependent size differences increased between groups. Larvae reared at the highest density (15-20 larvae/mL) exhibited reduced growth relative to lower density treatments, and consistently comprised the smallest size class from ∼30 DPF onward. In contrast, larvae reared at the two lowest densities (1 larvae/mL and 2 larvae/mL) attained the overall largest mean sizes throughout the measurement period. The intermediate density (5 larvae/mL) showed a growth trajectory that was intermediate between the lowest and highest densities (Figure 1).

**Fig. 1.**
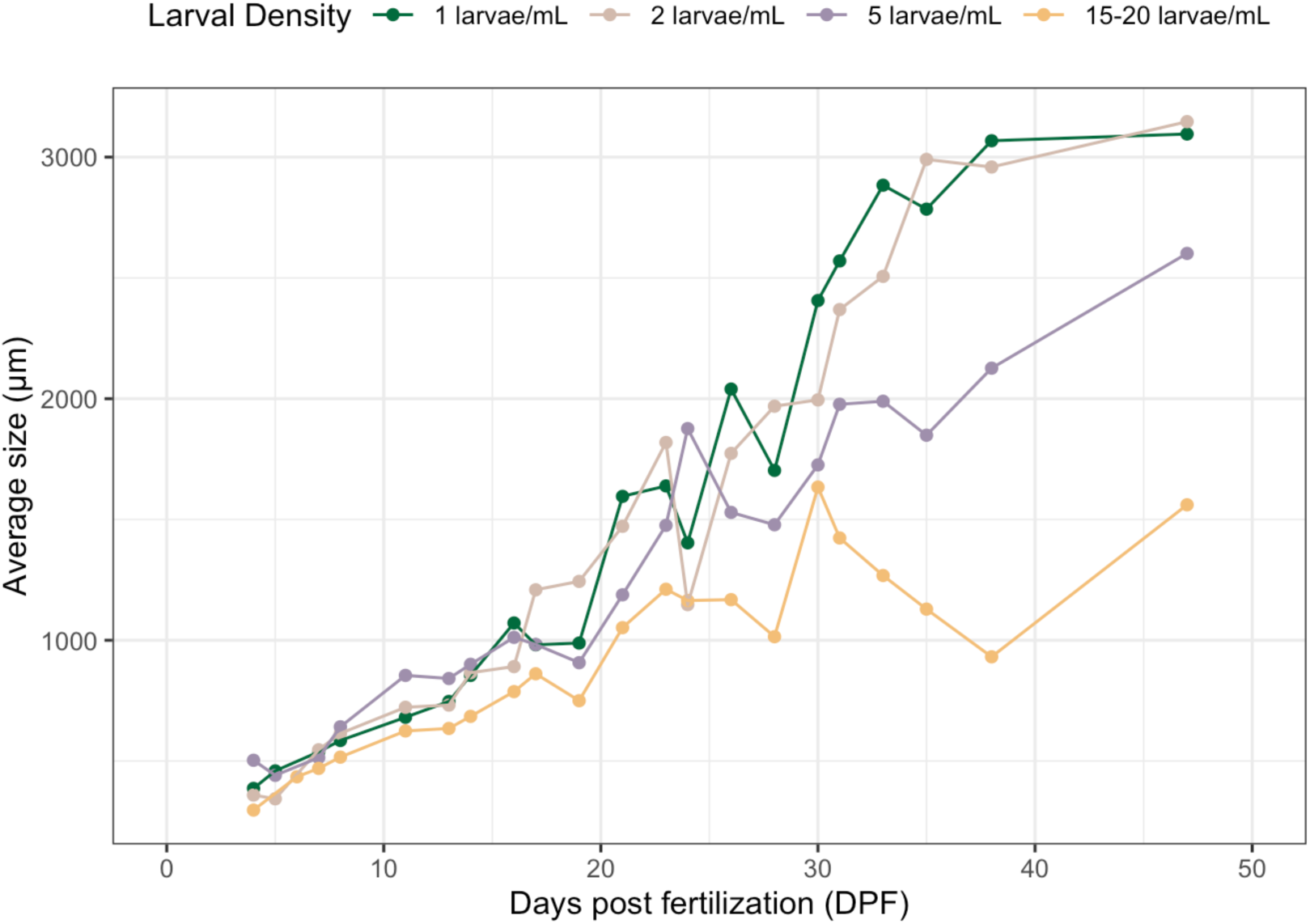
Larval growth trajectories according to larval density, measured from 4 to 47 DPF. Mean larval size (µm) is plotted over days post fertilization (DPF), with individual points representing means on the day of measurement for sub-sampled larvae from each density treatment.

**Fig. 2.**
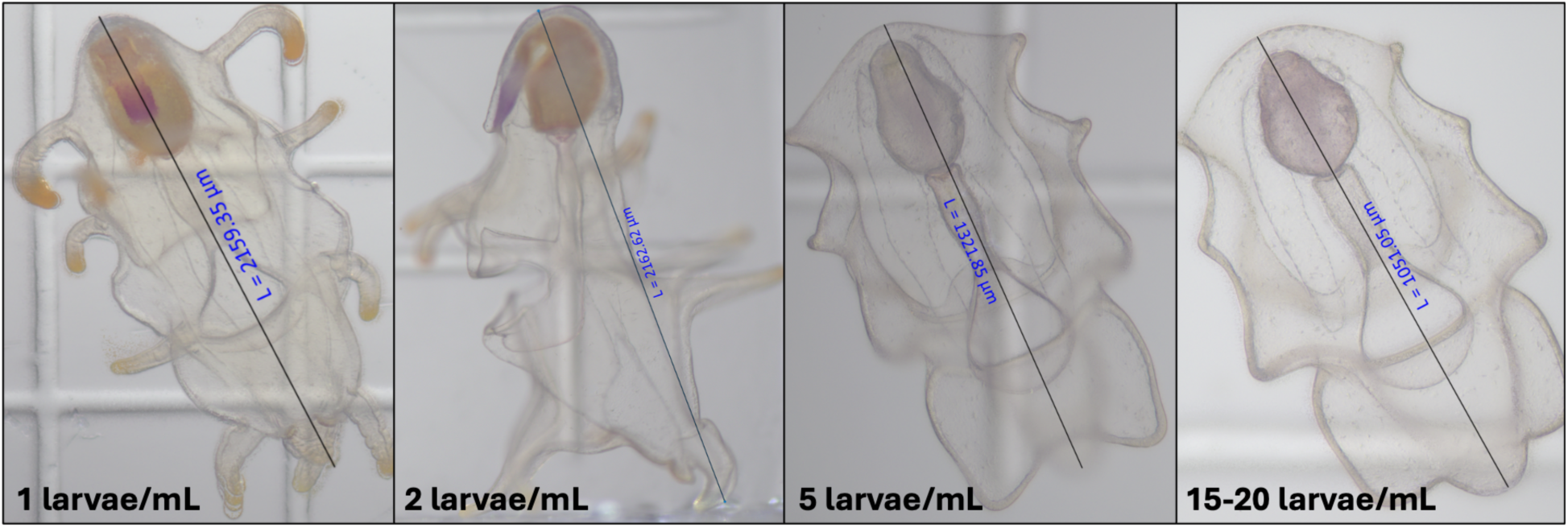
Photos of larvae captured 28 DPF, labeled according to density treatment each individual was sub-sampled from. Differences between stage-specific developmental features are most pronounced between the two lowest (left; 1 larvae/mL and 2 larvae/mL) densities when compared to the two highest densities (right; 5 larvae/mL and 15-20 larvae/mL).

Microscope photos captured throughout larval sampling showed differences in larval development stages between the two lowest and two highest larval densities at 28 DPF (Figure 2). The larvae subsampled from the two lower densities show larvae in the early brachiolarial stage, with brachiolar arm development clear in both photos. The larvae subsampled from the two highest densities were still in an earlier stage of bipinnarial development.

The first settled larvae were observed at 41 DPF and settlement numbers were recorded until 146 DPF. Cumulative larval settlement during the settlement period (3/26/2024 to 7/9/2024) increased across all density treatments (Figure 3A). The intermediate density, 5 larvae/mL, showed an early increase in settlement, but was later outpaced by the 15-20 larvae/mL treatment. Overall, end cumulative settlement differed significantly among density treatments (Figure 3B). When normalized by the number of replicates per treatment, the two highest densities yielded significantly greater numbers of settlers compared to the lowest density (One-way ANOVA, F(3,16) = 10.65, p <0.01; Tukey HSD, p < 0.05). End total mean cumulative settlement increased according to larval density (± SD): 1 larvae/mL (131.19 ± 84, *n* = 8), 2 larvae/mL (254.19 ± 116.69, *n* = 4), 5 larvae/mL (345 ± 107.85, *n* = 4), and 15-20 larvae/mL (448.31 ± 93.15, *n* = 4). At the end of the settlement period, the two highest densities yielded the largest number of cumulative settlers (normalized) compared to the lowest density (Tukey test, *P* < 0.01).

**Fig. 3.**
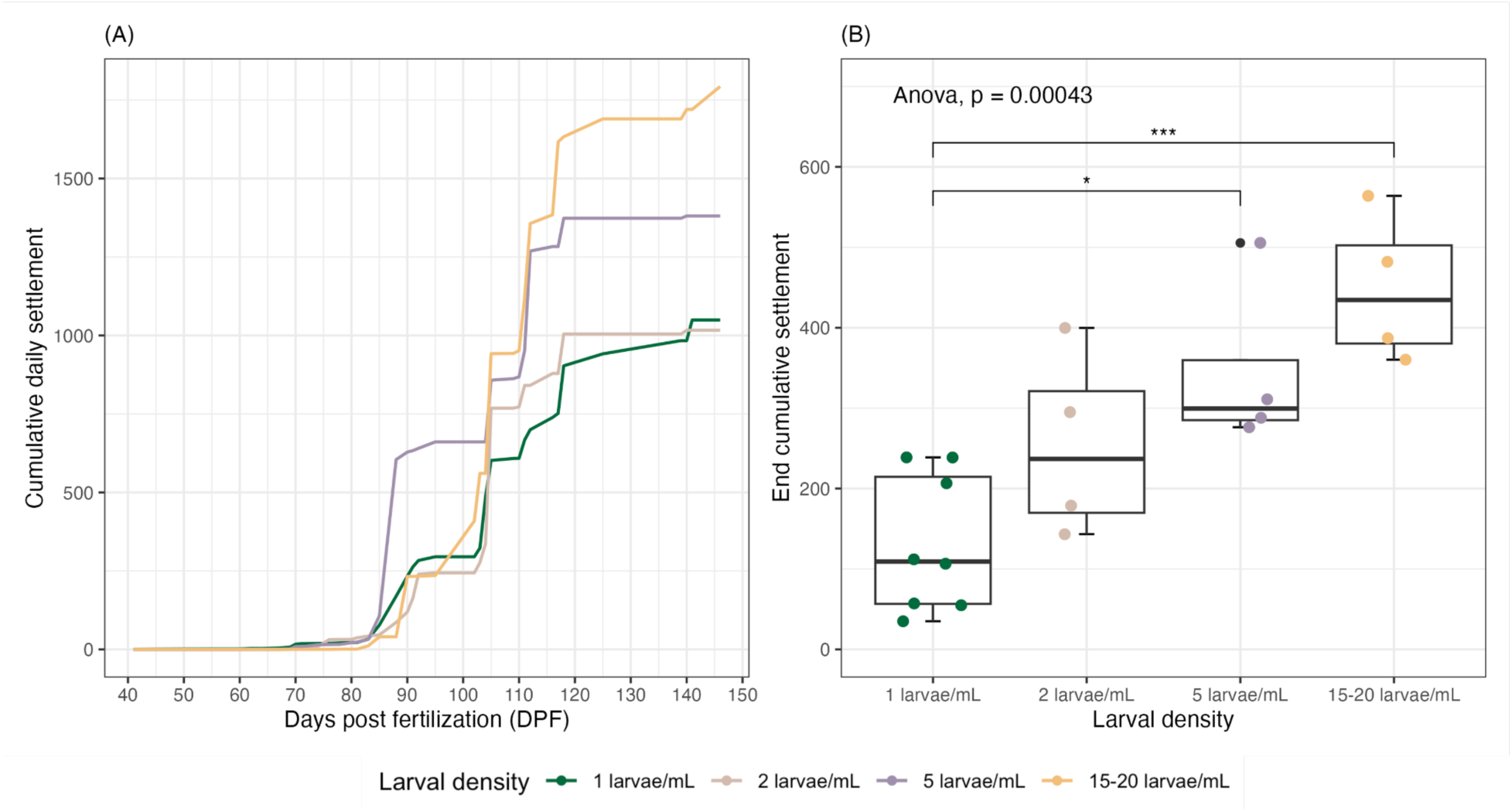
**(A)** Normalized cumulative daily settlement plotted over days post fertilization (DPF). Lines represent the progression of settlement for each larval density across the experimental period. **(B)** Normalized end total cumulative settlement of each larval density treatment 146 DPF. Individual replicates are overlaid as jittered points. Significance levels are indicated above the bars derived from the adjusted p-value from pairwise comparisons analysis (*P* < 0.05 (*), *P* < 0.01 (**), *P* < 0.001 (***)).

To examine settlement dynamics over time, we fit a generalized additive model (GAM) with a negative binomial distribution to normalized daily settlers as a function of DPF, allowing separate smooth terms for each larval density (Supplementary Figure 2). The parametric effect of density was not significant (*P* > 0.05), indicating no statistically significant shifts in the overall mean normalized settlement rate among density treatments. In contrast, smooth terms for DPF were significant for all densities (*P* < 0.001), showing strong, non-linear settlement dynamics over DPF within each density treatment (edf = 3.13-3.63). The model explained 31.1% of the deviance (adjusted R^2^ = 0.0648; *n* = 760).

The proportion of larvae that successfully metamorphosed and settled declined with increasing larval density (Figure 4). Mean settlement rate decreased with increasing larval density (± SD) 1 larvae/mL (13.2% ± 8.48, *n* = 8), 2 larvae/mL (6.35% ±, *n* = 4), 5 larvae/mL (3.45% ± 1.08, *n* = 4), and 15-20 larvae/mL (0.68% ± 1.14, *n* = 4). Settlement rate differed significantly among treatments (Kruskal-Wallis X^2^ (3) = 14.2, *P* = 0.0026), with the lowest density exhibiting significantly higher survivorship to settlement than the highest density (Dunn test, adjusted *P* < 0.05). Differences in the settlement rates between the lowest and the two intermediate density treatments were insignificant (Dunn test, adjusted *P* > 0.05).

**Fig. 4.**
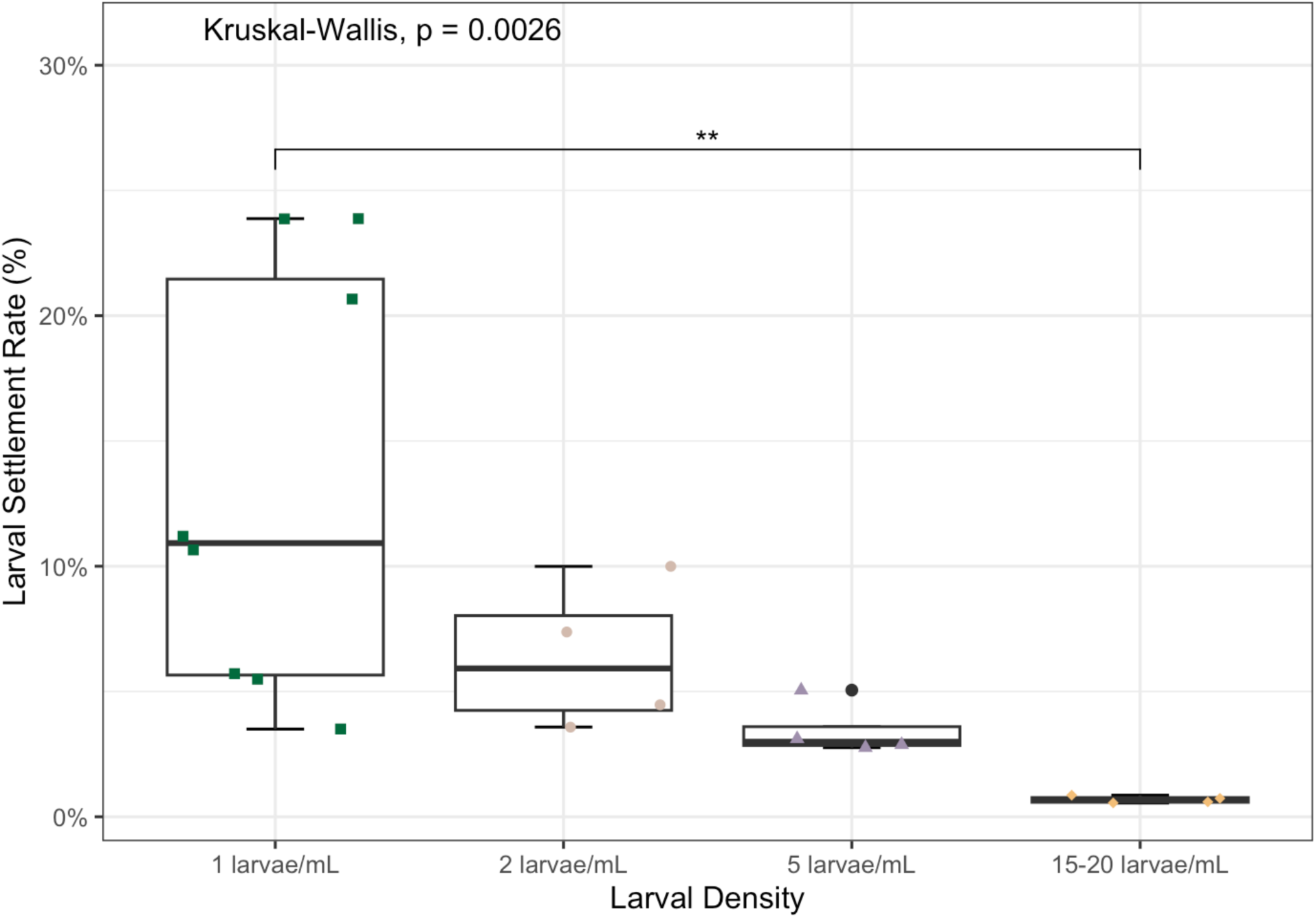
Percent settlement rate across the four larval density treatments. Individual replicates are overlaid as jittered points over boxplots. Between group significance levels are indicated above the bars as the adjusted p-value (*P* < 0.05 (*), *P* < 0.01 (**), and *P* < 0.001 (***)).

### One-year post fertilization size, survival, and fitness test by density

Overall, 102 juvenile Sunflower sea stars from the various jars survived to one year post-fertilization. The absolute number of surviving juveniles varied across treatments (1 larvae/mL, *n* = 31; 2 larvae/mL *n* = 12; 5 larvae/mL *n* = 26; 15-20 larvae/mL, *n* = 23) with an additional *n* = 13 whose original larval density was unknown, not tracked, or no measurements were recorded during the February 2025 fitness tests. Survival was calculated for each replicate as the proportion of juveniles recorded during the February 2025 fitness test relative to the number of settlers counted on July 9, 2024, the last day of settlement. Mean survival (± SD) from settlement to one year post-fertilization was highest for settled stars that had been reared at 1 larvae/mL (0.65 ± 0.56%) and lowest for those reared at 15-20 larvae/mL (0.32 ± 0.06%) (Figure 5). Mean survival was intermediate for jars reared at 2 larvae/mL (0.5 ± 0.18%) and 5 larvae/mL (0.44 ± 0.17%). The difference in means across larval treatments was not statistically significant (one-way ANOVA, F (3,16) = 0.706, *P* = 0.562).

**Fig. 5.**
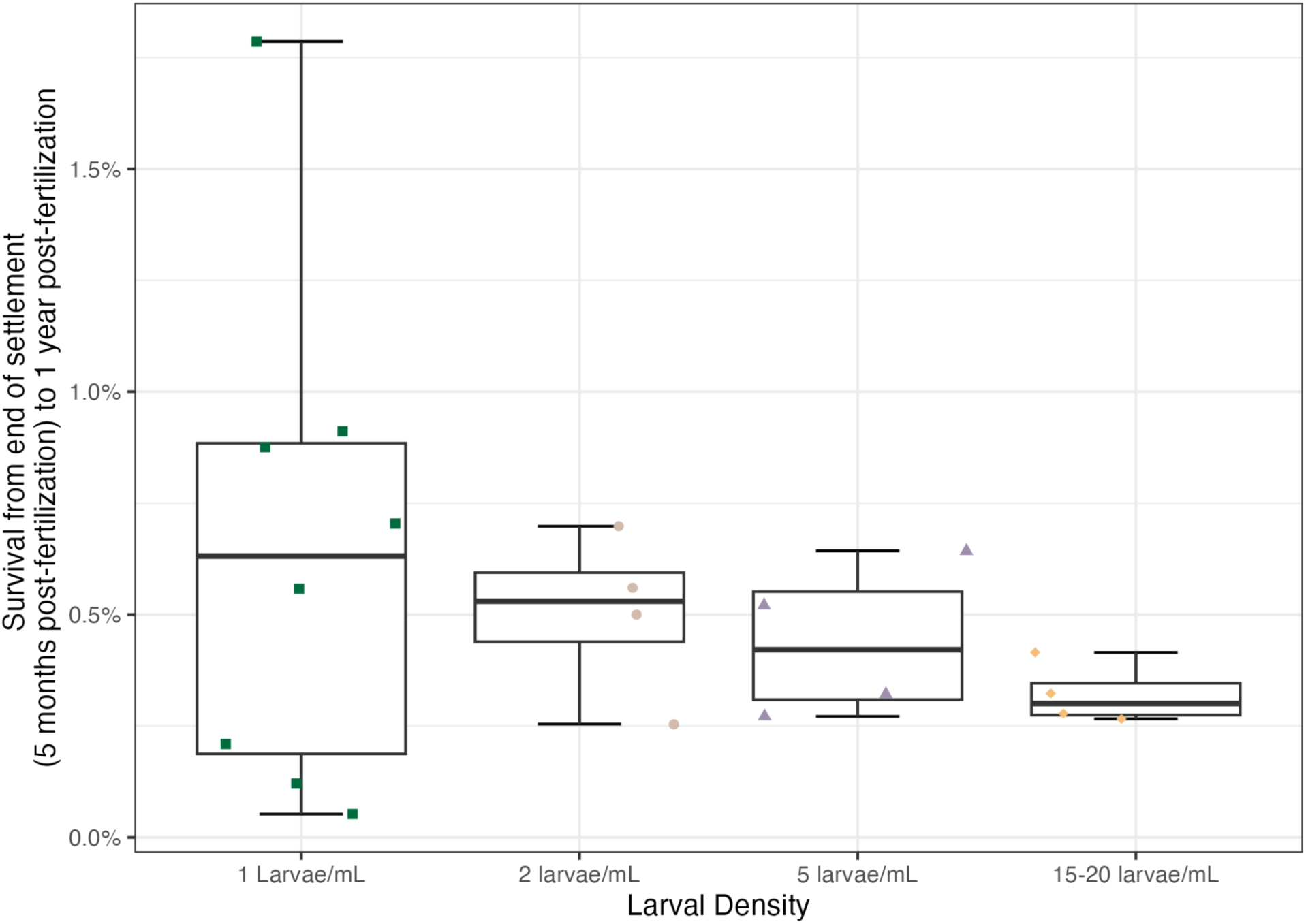
Juvenile survival from settlement to one year post-fertilization across larval density treatments. Density replicates overlaid as jittered points over boxplots.

Juvenile length at one-year post-fertilization ranged from 12 to 70 mm, and mean juvenile length was significantly different between larval rearing densities one-year post-fertilization (One-way ANOVA, F (3,84) = 7.72, *P* < 0.01) (Supp Fig 2A). Mean length (± SD) was largest in juveniles reared at the 1 larvae/mL (47.2 ± 11 mm) when compared to juveniles reared at 15-20 larvae/mL (41.7 ± 15.2 mm) and 5 larvae/mL (44.6 ± 11.7 mm) (Tukey test, *P* < 0.01) (Figure 6A). Patterns in juvenile weight (g) were similar to those observed in length, and after exclusion of one outlier, there was also a statistically significant relationship between juvenile weight and larval rearing density one-year post-fertilization (Kruskal-Wallis, X^2^ = 4.8, df = 3, *P* = 0.002) (Figure 6B). Mean weight (± SD) was significantly greater in juveniles reared at 1 larvae/mL (3.98 ± 2.38 g) and 5 larvae/mL (3.92 ± 2.54 g) when compared to the juveniles reared at 15-20 larvae/mL (1.88 ± 1.52 g) (Dunn test, all p < 0.01). There was no statistically significant difference in the number of arms a juvenile star had one-year post-fertilization among those reared at different larval densities (Kruskal-Wallis, X^2^ = 5.77, df = 3, p = 0.122).

**Fig. 6.**
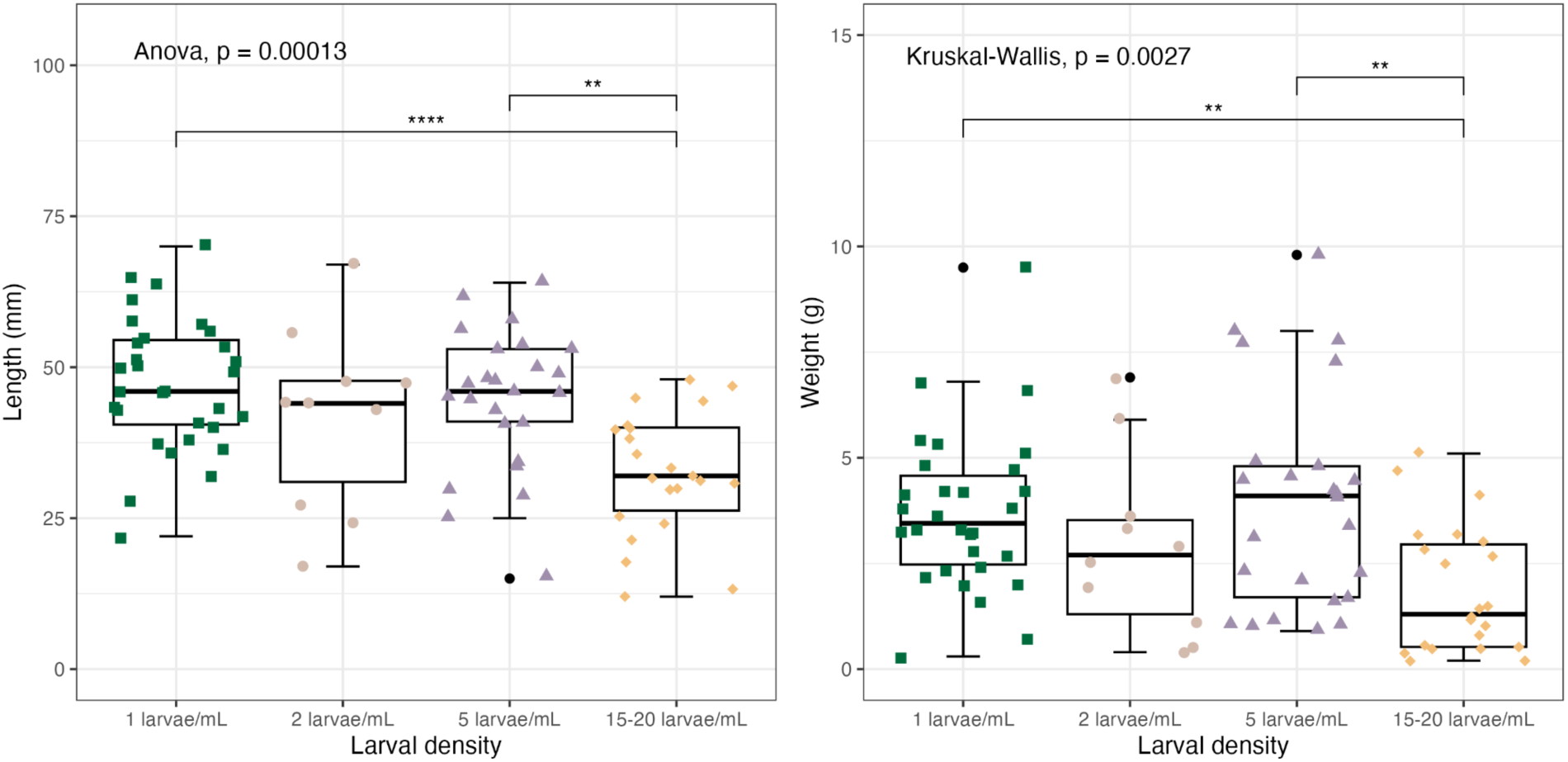
Significance levels are indicated above the bars as the adjusted p-value from pairwise comparisons (*P* < 0.05 (*), *P* < 0.01 (**), *P* < 0.001 (***), and *P* < 0.0001 (****)) (A) Juvenile armtip-to-armtip length (mm) one-year post-fertilization among larval density treatments. (B) Juvenile weight (g) one-year post-fertilization across larval density treatments.

We evaluated the relationship between body length and larval rearing density and flip time. Overall flip time showed substantial variability across individuals, with only modest amounts of variation explained by length and larval density (Figure 7). The mean flip times by larval density treatment were: 1 larvae/mL (415 ± 448 seconds), 2 larvae/mL (364 ± 351 seconds), 5 larvae/mL (279 ± 273 seconds), and 15-20 larvae/mL (443 ± 185 seconds). There was no significant difference in juvenile flip time among the larval density treatments (Kruskal-Wallis, X^2^ = 3.70, df = 3, p = 0.296) and pairwise comparisons detected no significant effect between density groups (Dunn test, P > 0.05) (Figure 7A).

**Fig. 7.**
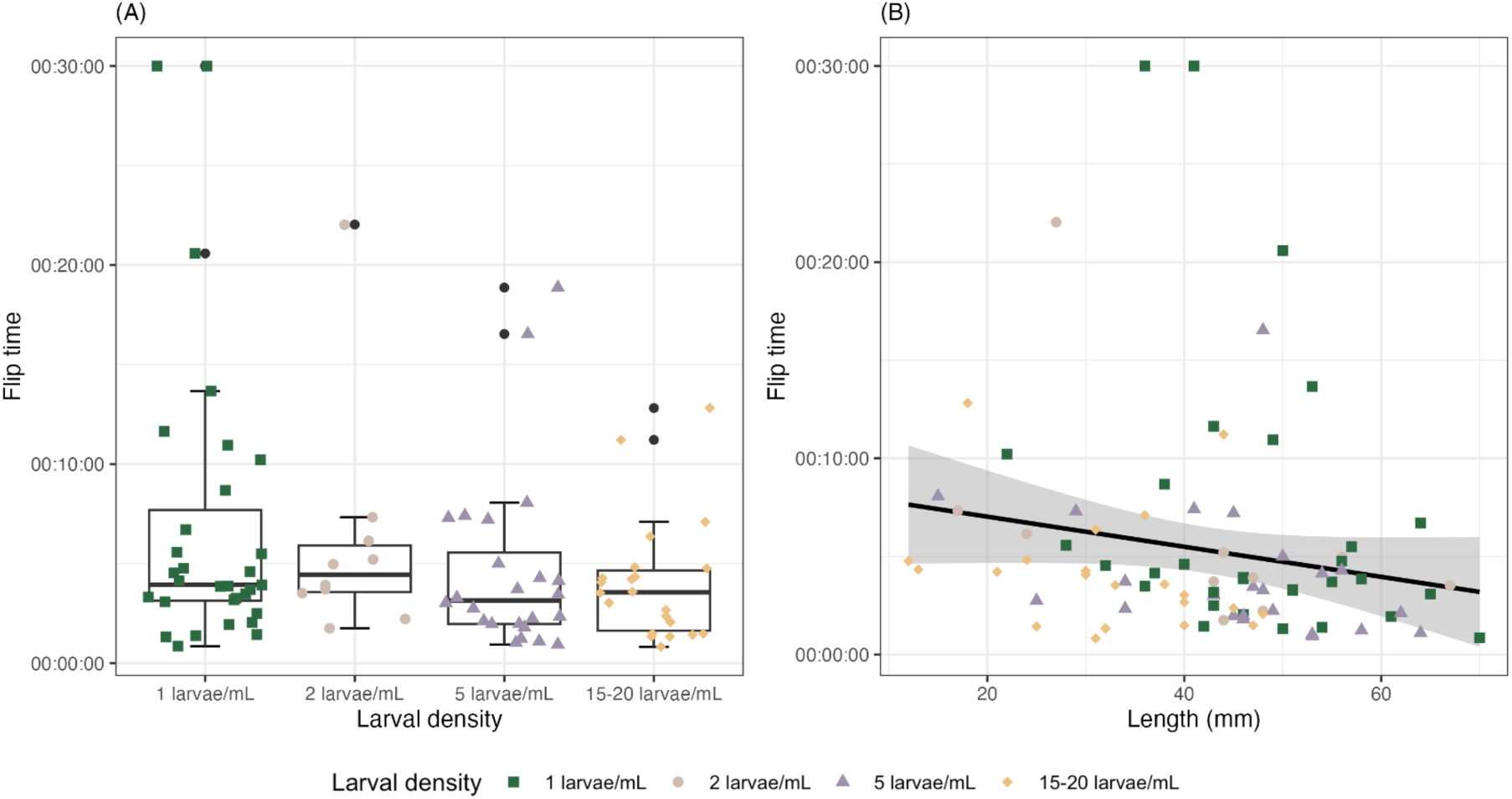
**(A)** Flip time(s) across larval density treatments. The vertical axis displays flip time as hours:minutes:seconds. **(B)** Relationship between length (mm) and flip time among larval density treatments. Points represent individual juveniles with shape and color indicating density treatment, reflected in the top legend. The solid black line shows a simple linear regression of change in flip time according to length, with the gray shading reflecting the standard error (± SE) around the fitted line.

Linear regression models explained very little of the variation observed in juvenile flip time. A simple linear regression indicated a weak, negative relationship between length and flip time, where larger bodied juveniles tended to flip slower than the smaller bodied juveniles (Figure 7B). This effect was marginal and not statistically significant (β = −4.6 seconds/mm ± 2.76 SE; F(1,84)= 2.76, *P* = 0.1) and length only explains 3.2% of the variation in flip time (R^2^ = 0.0318). There is a weak, negative relationship when allowing the effect of length to vary among density treatments (β = −4.176 seconds/mm:Density ± 2.33 SE; F (3,82) = 2.5, *P* = 0.066). This weak, negative relationship is not significant and the density interaction does significantly improve model fit or predictions compared to the simple model (F (2,84) = 2.3, *P* = 0.1). The addition of the interacting parameter explains 5% of the additional variation in flip time (R^2^ = 0.084, adjusted R^2^ = 0.05). The low amount of explained variance between the models leaves unexplained individual variation in the flip time performance.

### Relationship between length and arms, length and weight

We measured the length, weight, and arm count for every Cupid Cohort star included in the fitness tests at one-year post fertilization, as well as the length, weight, and arm count for 45 adult sunflower sea stars housed at the Seward Sea Life Center in Seward, Alaska. We also measured the length and arm count of 37 historical specimens held in the CAS Invertebrate Zoology collection, collected across the North American West Coast from 1889-2023. Weight of the historical specimens is uninformative, as these were either dry or preserved in ethanol, and as such their weights do not correspond to what the weight of the animal would have been while alive and therefore were not included in the weight analysis or modelling.

Across all specimens, body length was strongly associated with both arm count and body weight, with clear nonlinear relationships throughout development and growth (Figure 8). Arm count increased rapidly with length (cm) at smaller sizes and approached an asymptote in larger bodied individuals (Figure 8A). Fitting a self-starting nonlinear asymptotic model (SSasymp) to data from museum specimens, Alaskan stars, and the Cupid Cohort revealed that arm count likely asymptotes around approximately 19 arms (Asyms), with an initial arm count parameter of 3.41 (R_0_), and a rate constant of −1.91 (lrc). All of the model parameters were significant (*P* < 0.01), and overall model fit was strong (psuedo-R^2^ = 0.881). Bootstrapping (1,000 resamples) produced narrow confidence intervals around the mean prediction of arm count, indicating solid model-fit support for the asymptotic pattern. Observed arm counts ranged from a minimum of 7 arms from a one-year-old juvenile in the Cupid Cohort to 23 arms recorded in an adult Alaskan star. Arm counts across specimen types converged towards our modeled asymptote by approximately 20 cm in length.

**Fig. 8.**
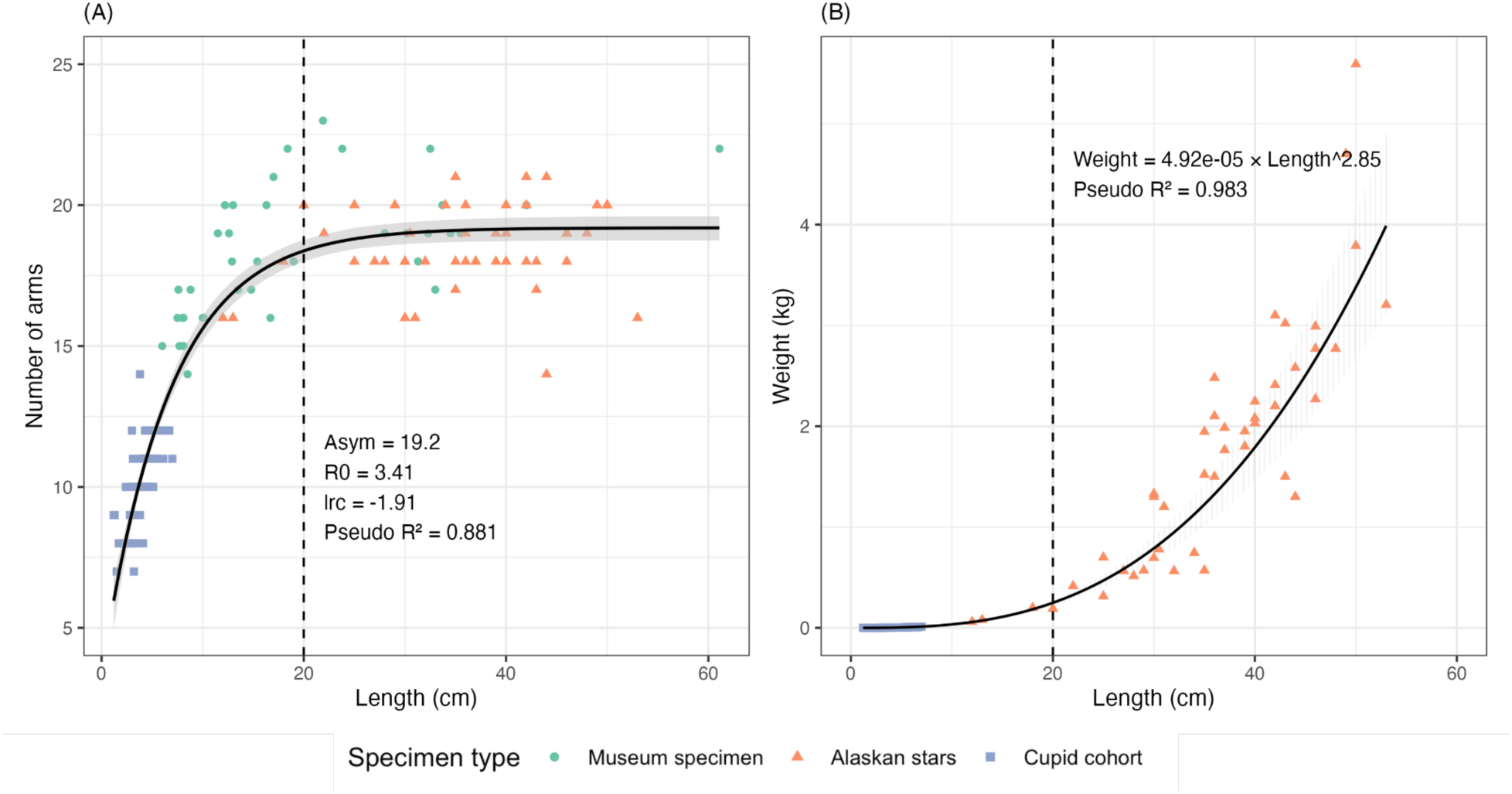
**(A)** Relationship between length (cm) and arm count of *Pycnopodia* across specimen types (museum specimens, Alaskan stars, and the Cupid Cohort). Points show individual observations, the solid black line represents the self-starting, fitted nonlinear asymptotic model (SSasymp), with a grey 95% bootstrap confidence interval (1,000 resamples). **(B)** Length-weight relationship for live stars. The solid black line is the fitted allometric model, with grey lines showing 1,000 bootstrap derived curves (95% CI). The vertical dashed line on both graphs marks the approximate transition where arm count asymptotes (A) and mass accumulation accelerates (B).

Weight (kg) recorded from the live sunflower sea star specimens contributed to increases in length (cm), following a power-law relationship as relative change in length results in proportional change in weight (Figure 8B). A log-linear regression model examining the potential of allometric growth yielded the model Weight = 4.92 x 10^-5^ x Length^2.85^, with the scaling exponent <3 (2.85), indicating negative allometry. Model fit was high (psuedo-R^2^ = 0.983).

Bootstrapped estimates of the scaling exponent were constrained (95% CI: 2.77-2.93), showing low uncertainty in our modeled relationship between the length-weight relationship across our measured size range from our various specimens. Weight also increases exponentially with the number of arms (Supplementary figure 3).

When visualizing the approximate length at which the number of arms asymptotes (∼20cm; Figure 8A), this also reflects the approximate point where weight begins to increase exponentially (∼20cm; Figure 8B). This is drawn as a vertical dashed line on both panels to reflect the hypothesized change in metabolic resource allocation from limb development to weight increase during growth.

## Discussion

The results of this work demonstrate success in spawn induction with 1-MeAde, larval rearing (including at much higher than previously reported densities; previous sunflower sea star larval rearing efforts have been done at 0.25 to 0.5 larva/ml [12]), and the size and fitness outcomes for juvenile stars one-year post-fertilization. From this, we can make recommendations about rearing larval and juvenile sunflower sea stars, and point to future areas of investigation.

### Larval Rearing

The biggest challenges faced in larval rearing were the time and labor required for regular water changes, and sufficient space to store larval cultures. Larval size was approximately similar across the four density treatments for the first 30 days, but after that the 1 and 2 larvae/ml treatments diverged and became larger on average than the 5 larvae/ml and 15-20 larvae/ml treatments. Images taken at 28 DPF indicated that development was faster in the lower density treatments, with those larvae shifting into the brachiolarial stage, while the higher density treatment larvae stayed in the bipinnarial stage longer. These results indicate early divergence in density dependent growth and development between the lowest and highest larval densities, also reflected in the size measurements (Figures 2 and 3). Larval size measurements stopped being recorded at day 47, shortly after larvae started settling (41 DPF). At this time, the 1 and 2 larvae/ml larvae were approximately 20% larger than the 5 larvae/ml treatment and about 100% larger than the 15-20 larvae/ml treatment. Peak settlement time was roughly the same across all four density treatments based on the results of the GAM, so it appears that the higher density larvae caught up in development eventually (Supplementary Figure 2).

A key feature in the way the larvae were fed in this experiment is that all cultures were fed to satiation. Because all cultures were fed until all the larvae in the culture had eaten, any differences in size or settlement rate across the different treatments are less likely to be a result of increased competition for food in the higher density larval cultures, and more likely to be due to space constraints or water quality differences. This is different from most previous studies, with the exception of [30], who similarly fed their different density treatments of purple sea urchin (*Strongylocentrotus purpuratus*) to satiation as determined by gut content checks, and similarly found faster larval growth in their 0.5 and 1 larvae/ml cultures than their 2 or 4 larvae/ml cultures. Metabolic waste accumulation and competition for space seem to be the limiting factors for growth in these cases where competition for food is not the limiting factor [30].

The total number of settlers was highest in the highest density treatment, but the rate of settlement, defined as the proportion of stars that settled out of the total starting number of larvae in a jar, was inversely proportional to larval density. A similar pattern has been found in the tropical crown-of-thorns sea star *Acanthaster cf. solaris*, in which the settlement rate for larvae kept at 2 larvae/ml was an order of magnitude lower than those kept at 1 and 0.5 larvae/ml [17]. The highest larval culture conditions in that study were the same density as the lowest density culture conditions in our study. That we were able to produce settlers at even higher densities is likely attributable to the fact that we fed our star larvae to satiation, as opposed to keeping food density constant across all cultures. This suggests that competition for food is a greater determinant of larval settlement than larval density itself. Interestingly, in a study of the gastropod *Babylonia formosae*, which has lecithotrophic (non-feeding) larvae, an opposite pattern has been found, in which rate of settlement increased with increasing density [31]. Those authors hypothesized that lecithotrophic larvae accelerate their development to reduce the amount of time spent in the dense, suboptimal larval environment. In contrast, feeding larvae like those of the sunflower sea star seem to experience delayed or arrested development when larval concentration becomes very high. While cumulative settlement in the two highest densities may have out-paced the lowest density, conservation breeding programs have to contend with the trade-off between the absolute number of settlers and rate of settlement when picking the density at which to raise their larvae. One other consideration currently under investigation is whether the genetic composition of the settlers changes when the rate of settlement decreases due to increased larval density.

### Juvenile rearing and fitness

There were no significant differences in juvenile post-settlement survival from the end of the settlement period to one-year post-fertilization. However, the highest density treatment led to significantly smaller juvenile stars one year after fertilization. Previous studies on other marine invertebrates have shown that stressful larval experiences can lead to reduced growth rate at the juvenile stage (e.g. in shore crabs [32] and in Olympia oysters [19]).

Flip time is routinely used to measure adult echinoderm fitness, and has more recently been applied to juveniles [13]. There was no statistically significant difference in flip time for any density treatment, and there was very little to no correlation between size (measured in terms of length or weight) and flip time generally. This suggests that, while the juveniles raised at the highest density as larvae may have experienced some energetic deficits leading to reduced size, the size difference does not indicate a reduction in fitness at one year old. This contrasts with findings from juvenile crown-of-thorns sea stars, for which flip time was faster in larger juveniles [33].

The biggest challenges in rearing juvenile stars included providing sufficient food, cannibalism, space, cleaning, and pest outbreaks. Limited staffing and the large amount of time needed for cleaning resulted in breakouts of benthic copepods that restricted growth of newly settled stars. Copepods were never directly observed feeding on newly settled stars, but bins with large outbreaks experienced high mortality. These copepods were observed hunting and eating newly settled urchins, so it may have been that they were outcompeting the sunflower sea stars for prey items.

### Allometric growth

When comparing the length, weight, and number of arms across the juveniles raised in this study with live adult animals collected in the ocean, and comparing the length and arm counts across live animals and preserved historical animals, patterns of allometric growth emerge. At smaller sizes, growth in the number of arms increases rapidly, but then begins to level off around the point the animal becomes 20 cm in length. Also at approximately 20 cm in length, the rate of increase in weight really begins to accelerate. This significant exponential relationship between diameter and weight has also been observed in juvenile crown-of-thorns sea stars [34]. The energetic cost of growing an arm is very high for echinoderms [35]. We found a similar exponential relationship between number of arms and body weight in sunflower sea stars as was found by [36] (Supplementary Figure 3). McClintock (1989) also found that as sunflower sea stars grow from juvenile to adult size, the protein and lipid content of the body wall increases, and the energy content of the body wall tissues increase 26%. McClintock suggested that the change in energy content allows the adult tissues to serve as reservoirs for gametogenesis. From this, we hypothesize that the sea stars’ metabolic allocation shifts from limb development to body mass as they grow and mature, in order to reach reproductive maturity and successfully produce gametes. In practical terms for aquarium biologists and other husbandry practitioners, it is useful to know when to expect weight gain, and demand for food, to accelerate. This may also aid in understanding tank size demands as juveniles grow.

### Conclusions

We show that successful sunflower sea star rearing can be achieved, even at much higher than previously reported larval densities. While larval density influenced larval size and settlement rate, it ultimately did not affect the survival rate or fitness rate post settlement, and affected juvenile size only for the highest density group. Decisions about density can be made based on the constraints of the larval program’s space, without too much concern that a 1, 2, or 5 larvae/ml density will adversely affect the outcomes measured here. However, changing density may affect the genetic composition of the surviving stars-this is currently under investigation.

We can also share anecdotally that denser cultures are harder to keep clean, due to the amount of waste products produced, and therefore require many more person-hours to maintain.

Therefore, we recommend keeping cultures at 1-2 larvae/ml to reduce husbandry staff time requirements for this species, and for other feeding marine invertebrate larvae with long larval duration times.

## Data Availability

All data used for analysis are available in the form of .csv files in the Supplementary Tables section. All code used for analysis is available in the Rmarkdown file in the Electronic Supplementary Material, and is publicly available at https://github.com/tallulahstory/Pycnopodia-aquarium-husbandry---Cupid-Cohort.

## Acknowledgements

Funding for this project came from Bill Younger, the Gardner family, the National Science Foundation Research Experience for Undergraduates program (Award Number 2243994), California state earmark GF2226-0, and National Oceanic and Atmospheric Administration grant (Federal Award Identification Number: NA24NMFX463C0055-T1-01 / NA24NMFX463C0055 / Mod 0). We thank our collaborative partners that made the Cupid spawn possible: Aquarium of the Pacific, Birch Aquarium, San Diego Wildlife Alliance, and Sunflower Star Laboratory. We thank Johanna Loacker for her assistance with CAS Invertebrate Zoology specimen acquisition for historical measurements. We thank Hog Island Oyster Co. for providing some of the live food items for our juvenile sunflower sea stars. Finally, we thank the Animal Care, Animal Health, and Water Quality teams at the Steinhart Aquarium for their passion and dedication to the care of these animals, and all of the animals, at the aquarium each and every day.

