## Supplementary Figures for "Growth, survival, and fitness in the first year of life for *Pycnopodia helianthoides* under different larval densities"

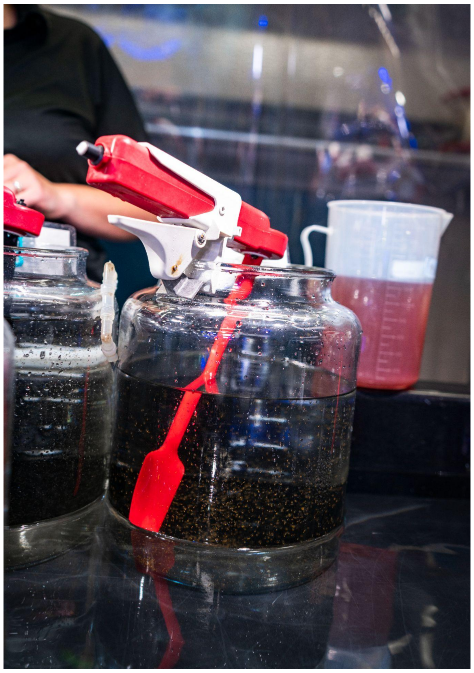

**Supp. Fig. 1.** We reared larvae in 8 liters of water, using a StirMATE modified with a spatula attachment to produce water movement.

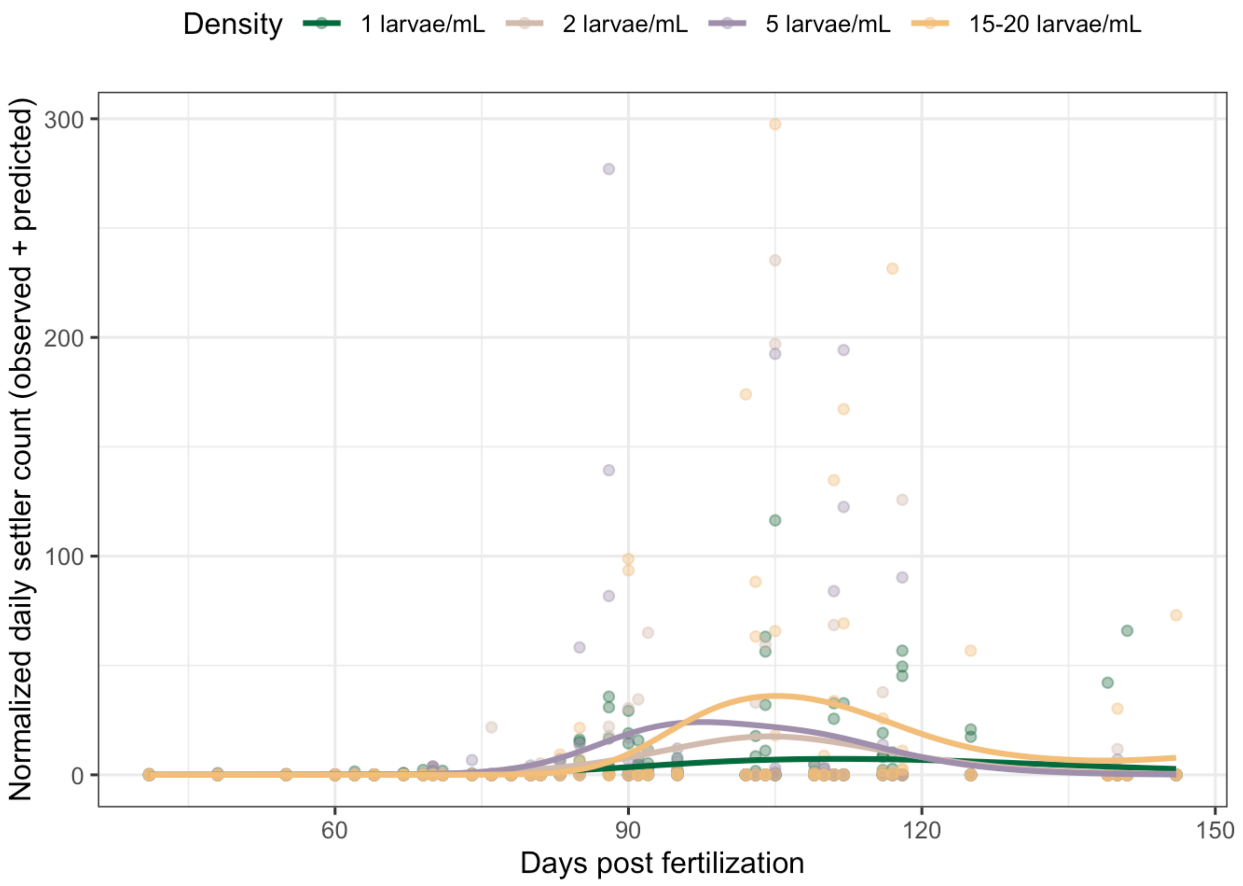


**Supp. Fig. 2.** Observed and generalized additive model (GAM)-predicted of daily larval settlement as a function of days post fertilization (DPF) across the four larval rearing densities, shown in top legend. Points represent the observed values and lines represent the GAM predictions.

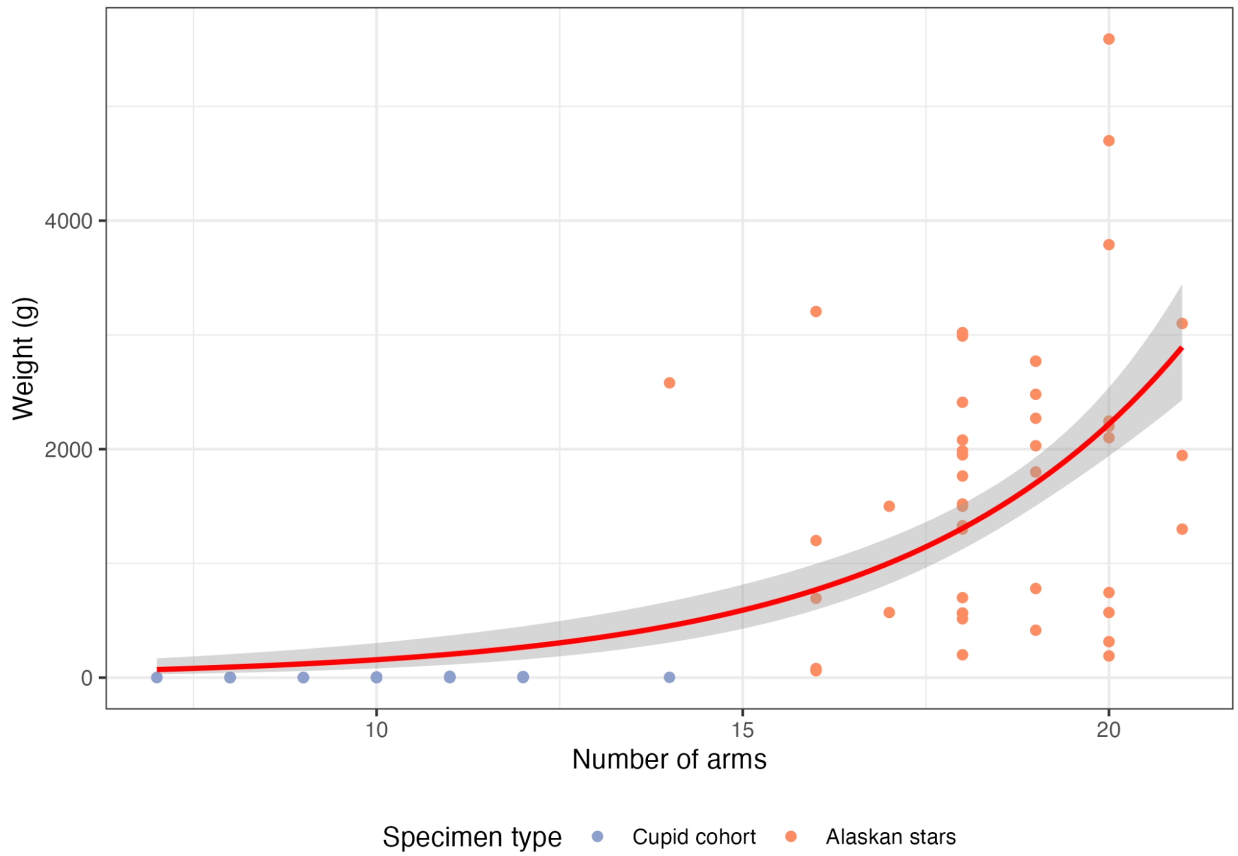


**Supp. Fig. 3.** Relationship between arm count and weight (g) across juvenile and adult sunflower sea star specimens (Alaskan stars and Cupid Cohort). Points show individual observations. The red line shows a fitted gaussian GAM smooth, with the grey shadowing around the line showing the model SE.
