## Supplementary Methods for "Growth, survival, and fitness in the first year of life for *Pycnopodia helianthoides* under different larval densities"

1. **Spawn Induction**
2. **Aquatic live food cultures- phytoplankton**
1. **SPAWN INDUCTION**

To prepare a 75 ppm (75 mg/L) solution of 1-MeAde, we mixed 25 mg of 1-MeAde with 334 mL of 1 micron filtered seawater. The solution was heated to 30 °C while stirring until the powder was fully dissolved. Once the powder was fully dissolved, we cooled the solution to the same temperature as the water that the target animals were kept in, in order to avoid heat shock when injected.

We weighed each sea star, and then drew up the dose of 1-MeAde at 1 mL per 150 g body weight. We then rinsed the desired injection site copiously with sterile saline that was similar in temperature to the water that the target animals were kept in. We administered the injection using a sterile, new 20 gauge needle attached to an IV extension set. We positioned the needle at a 45-90°angle to allow for easier entry through the epidermis and calcium carbonate endoskeleton. The solution was injected intracoelomically in the dorsal (abaxial) surface of an arm, midline, halfway from the central disc. For consistency of injection, the arm closest to the madreporite was chosen. We injected the solution at a slow, steady rate so that the full dose was given over 10 minutes, in order to minimize trauma to the animal. Each star was then monitored to determine the length of time between injection and spawning.

2. **AQUATIC LIVE FOOD CULTURES - PHYTOPLANKTON**

I. **System Specifications**
   A. Towers and Carboys located in Cultured Feed Room (CFR), Parent Stock rack located in B1E degas room
   B. Three 240-260 Liter conical bottom translucent towers containing T-*Isochrysis* and two 90-110 Liter conical bottom translucent towers containing *Rhodomonas*
   C. Up to nine 20 Liter carboys on carboy rack
   D. Parent stock consists of 2L flasks
   E. Air and CO_2_:
1. Building compressor aerates all cultures and it is injected with CO_2_ via a tee in the airline before reaching the cultures
2. Compressor air is filtered in-line with 45 micron, 5 micron, and 0.2 micron filters. Filters should be changed every 4 months or more often as needed.
3. Tower air injected into bottom of culture via tee
4. Carboy and flask air injected into top of culture via rigid airline tubing
5. CO_2_ regulator model Green Leaf Aquarium dual stage model 103101. A 20 pound CO_2_ bottle typically lasts up to 4-6 months.
6. Backup plug in air pump can be used if building air is not available via tee in airline and valve isolation; Pentair Sweetwater Linear II Diaphragm air pump model SL94A
7. Parent stock is only kept on air only without CO2 infusion via local Pentair sweetwater pump on a 0.2micron filter
   F. Lighting:
1. Towers: Twelve 48” fluorescent fixtures 4200K or LED lamps, vertically hung, 24 hour photoperiod, fluorescent lamps changed every 6 months, LED lamps changed yearly
2. Carboys: Six 24” fluorescent fixtures, 4200K or LED lamps, horizontally hung, 24 hour photoperiod, fluorescent lamps changed every 6 months, LED lamps changed yearly. Rhodomonas has only 1 fixture (2 lamps) operational due to lower light requirements.
3. Parent Stock: two 48” fluorescent fixtures 4200K, horizontally hung per shelf, 24 hour photoperiod, lamps changed every 6 months. Rhodomonas has only one lamp operational due to lower light requirements.
II. **Species List**
    A. Phytoplankton species
    1. T*-Isochrysis galbana*
    2. *Rhodomonas lens*
    3. *Chaetoceros gracillis*
    4. *Nannochloropsis* sp.
    5. *Tetraselmis* sp.
    6. *Nitzschia pusilla*
IV. **Maintenance**
    A. Daily Maintenance
       1. Check/record CO_2_ bottle pressure to verify gas delivery to phytoplankton cultures. Working pressure should be 32 psi and the bubble counter should be steady and approximately 2 bubbles per second
       2. Check air flow to all cultures. If air is low, increase airflow to culture and/or change 0.2 micron inline disc filter. If foam is forming at the surface of culture, it is likely air needs to be reduced- this is most prevalent in Rhodomonas cultures.
       3. Harvest daily general use amounts into general use vessels or pitchers from towers and placed in designated harvest area. Copepod and rotifer feed harvests should be added directly to the cultures, rocky reef and other specialty harvests will be set aside for the specific biologists. Check the whiteboard daily to see harvest amount updates.
       4. Parent stock should be swirled AM and PM and temperatures recorded. If temperatures fall outside of 70-80F take action to reduce or increase temperatures.
    B. Tower cleaning
       1. After the harvest tower is empty, turn off air completely and spray the tower down with freshwater. Rinse and wipe off place plastic tower dust covers and place in bleach bucket
       2. Keep harvesting valve open and plug hose with pvc plug
       3. Refill with freshwater to highest point possible, adding 500ml 8.25% bleach and 1L bleach for large towers, add bleach while filling to mix and ensure airline is bleached
       4. Let sit overnight (preferably) or a minimum of 3 hours, make sure there is no aeration that can remove bleach from water
       5. After bleaching is complete remove plug, drain and thoroughly spray clean with freshwater
    C. Carboy cleaning
       1. Spray carboy out with freshwater
       2. Refill with water to the base of the neck and top off with bleach to the highest point possible (approximately 250ml)
       3. Rigid airline and cap can be cleaned in bleach bucket overnight, be sure to rotate the rigid airline to ensure both sides come into contact with the bleach
       4. Next day, drain and rinse carboy thoroughly and thio/rinse airline and caps
       5. If not acid washing immediately, store airline and caps in the enclosed plastic bin above glassware and carboys on shelf
    D. Acid Washing and Inoculation Prep: This should be done at least the day before inoculation to allow bleach to fully disinfect saltwater
       1. All acid washing should be done with appropriate personal protection equipment: goggles and gloves
       2. Tower prep:
          a. Rinse down tower with dechlorinated freshwater ensure all bleach has been removed
          b. While the tower is still very wet, acid wash the tower with 10% hydrochloric acid. Use squirt bottle and steadily rinse tower down three times coating entire interior surface
          c. Use wypall to wipe top rim of tower and any water stains on outside of tower
          d. Let stand 10-15 minutes
          e. Rinse with dechlorinated freshwater thoroughly
          f. Close valve
          g. Change gloves
          h. Purge saltwater line and wipe down and douse candy cane and valve with 95% Ethyl Alcohol, allow alcohol to dry
          i. Purge saltwater line and allow saltwater to flow over candy cane removing any residual alcohol
          j. Fill with tower with saltwater to desired level
          k. Sterilize saltwater by adding 1ml 8.25% bleach per 10L of saltwater
          l. Place freshly bleached/naturalized/rinsed plastic dust cover on top of tower
          m. Let stand overnight. There should be no aeration which could remove bleach from the saltwater. Towers can have prepped bleach/saltwater for up to a week
       3. Carboy and flask Prep:
          a. Rinse carboy or flask with dechlorinated freshwater
          b. Add 10% hydrochloric acid to carboy and coat all sides 3 times and swirl, ensuring to coat the water line area thoroughly
          c. Rinse rigid airline tubing, glass tubing, caps and/or plugs with 10% hydrochloric acid 3 times
          d. Let stand 10-15 minutes
          e. Rinse carboy, flask, airline tubing and caps with dechlorinated freshwater thoroughly
          f. Change gloves
          g. Purge saltwater line and wipe down and douse candy cane and valve with 95% Ethyl Alcohol
          h. Fill with saltwater to desired levels: 5 gallon mark for carboys and 500ml or 2L for flasks
          i. Sterilize saltwater by adding 1ml 8.25% bleach per 10L of saltwater- line on carboy is close enough to 20L to add 2ml for sterilization, 500ml flask add 1 drop, 2L flask add 4 drops
          j. Add airline and cap to carboy or flask
          k. Cover airline and cap with parafilm or lid caps, breaking the parafilm over the airline to create a seal
          l. Let stand overnight at or 3 hours at a minimum, there should be no aeration. Carboys and flasks can have prepped bleach/saltwater in them for up to 1 week as long as the airlines and caps are sealed.
    E. Inoculation
       1. Change gloves and wipe down counter with Ethyl Alcohol
       2. For carboys and flasks, preferably place them next to cultures to be inoculated to match temperature overnight
       3. Neutralize chlorine in towers, carboy or flask with an equal amount of sodium thiosulfate solution (see solution preparation in notes) as bleach added- 1ml per 10L of saltwater
       4. Aerate towers and carboys vigorously for several hours or overnight to ensure bleach is neutralized. If in doubt a chlorine test can be conducted
       5. Return aeration levels to desired amount for phytoplankton growth, Rhodomonas should be slightly lower than all others
       6. Add fertilizer to towers and carboys and flasks
          a. Change gloves before opening any container with sterilized saltwater
          b. Fertilizer is 2 parts, Fritz Aquatic Pro F/2 part A and part B at the concentration of 3ml per 10L prepped saltwater. Note this is 3x more than the manufacturer recommended amount
          c. Shake fertilizers vigorously before use
          d. Add to towers, carboys, flasks, using clean graduated cylinders according to chart below

| Vessel | Part A | Part B |
| --- | --- | --- |
| 2L Flask | 1ml | 1ml |
| 20L Carboy | 6ml | 6ml |
| Towers | 3ml per 10L water | 3ml per 10L water |

e. Diatoms such as Chaetoceros and Nitzschia require sodium metasilicate to be added to the culture, 0.1g/10ml saltwater. This is lower than the recommended dosage due to the high silicate content in our saltwater.
7. With 95% Ethyl alcohol, wipe down neck and top portion of carboy or flask being used for inoculation to prevent any contamination
8. Pour desired amount of phytoplankton into freshly prepared tower, carboy or flask and replace covers immediately. If possible do not use any items like funnels that can be potential sources of contamination. All phyto is inoculated from flask to carboy to tower. Depending on current needs, inoculation can be light or dark. General inoculation volumes:
   a. 500ml Flask to flask inoculation 25ml
   b. 2L Flask inoculation is 100ml for rhodo, 250ml for iso
   c. Flask to carboy inoculation use entire 2L flask
   d. Carboy to tower inoculation use entire 20L carboy
9. If inoculating multiple species of algae or the same species from a different carboy, change gloves again and repeat
10. Generally, iso flasks should mature in 2-3 weeks, rhodo flasks in 1 week, carboys in 1-2 weeks and towers 3-5 days depending on inoculation density and volume.

V. **Notes**
A. Sodium thiosulfate solution is made by mixing 60g sodium thiosulfate granules in one liter of RO/DI water with the magnetic stirrer until completely mixed. This solution is a 1:1 bleach neutralization ratio specific to concentrated 8.25% bleach
B. If a culture appears contaminated by becoming light, cloudy, and/or has coagulated cells/chunks discard the culture and start it again. Flasks and carboys can be split multiple times at once at high or low density to replenish replicates
C. CO2 levels can be verified by taking the pH of a culture. Desired pH is 8-8.5, if pH is 9.5 or above more CO2 is needed
D. Notes on species:
   1. Rhodomonas does better with less light and cooler temps and if over-aerated foam will appear, usually with green cells in the foam. Placement at the end of a light strip or pulled further away from the light works well with this species. Keep on a slightly cooler shelf.
   2. Aeration and temperature of Iso, Tet, Nanno and Chaeto, Nitzschia can be identical
   3. Diatoms requires sodium metasilicate to be added at same time as fertilizer
   4. Cultures should be split before becoming overly mature, introducing dead cells into new cultures
E. Avoid cross contamination with other live food culture or exhibit work
   1. Always work with phytoplankton first
   2. Thoroughly clean hands and arms with soap and water when moving between live foods cultures or after working on any habitats
   3. If possible, space out doing different live food cultures over time, such as doing other cultures after lunch or later in the day
   4. If possible, different people should do different live foods culture care
   5. Order of culture work to reduce contamination: Phytoplankton parent stock->CFR Phytoplankton->Copepods->Rotifers->Artemia
